# Rare instances of haploid inducer DNA in potato dihaploids and ploidy-dependent genome instability

**DOI:** 10.1101/2020.11.02.365601

**Authors:** Kirk R Amundson, Benny Ordoñez, Monica Santayana, Mwaura Livingstone Nganga, Isabelle M Henry, Merideth Bonierbale, Awais Khan, Ek Han Tan, Luca Comai

**Affiliations:** Plant Biology and Genome Center University of California, Davis, 1 Shields Avenue, Davis, CA, 95616, USA; International Potato Center (CIP), P.O. Box 1558, Lima 15024, Peru; School of Biology and Ecology, University of Maine, Orono, ME 04469, USA; Current Address: Duquesa Business Centre, P.O. Box 157, Manilva, Malaga, Spain 29692; Current Address: Plant Pathology and Plant-Microbe Biology Section, Cornell University, 14456 Geneva, NY, USA; Integrative Genetics and Genomics Graduate Group, University of California, Davis, CA 95616, USA

## Abstract

In cultivated tetraploid potato, reduction to diploidy (dihaploidy) allows hybridization to diploid germplasm, introgression breeding, and may facilitate the production of inbreds. Pollination with haploid inducers yields maternal dihaploids, as well as triploid and tetraploid hybrids. It is not known if dihaploids result from parthenogenesis, entailing development of embryos from unfertilized eggs, or genome elimination, entailing missegregation and loss of paternal chromosomes. A sign of genome elimination is the occasional persistence of haploid inducer DNA in some of the dihaploids. We characterized the genomes of 1,001 putative dihaploids and 134 hybrids produced by pollinating tetraploid clones with three haploid inducers, IVP35, IVP101, and PL4. We detected inheritance of full or partial chromosomes from the haploid inducer parent in 0.87% of the overall dihaploid progeny, irrespective of the combination of parental genotypes. Chromosomal breaks commonly affected the paternal genome in the dihaploid and tetraploid progeny, but not in the triploid progeny. Residual haploid inducer DNA is consistent with genome elimination as the mechanism of haploid induction. Further, the fact that paternal chromosome breaks are specific to dihaploids and tetraploid progeny suggests that they may be specific to 2x sperms, and supports the hypothesis that 2x sperms facilitate genome elimination.

## Introduction

Cultivated potato (*Solanum tuberosum* L.) is predominantly autotetraploid (2n=4x=48), vegetatively propagated, highly heterozygous, and can be severely affected by inbreeding depression. These attributes make potato improvement through conventional breeding slow and difficult, and have renewed efforts to reinvent potato as a diploid and inbred-based crop based on true seed, in order to keep pace with a rapidly changing market and climate (Jansky et al., 2016). The first step is to capture useful genetic diversity of elite tetraploid cultivars at the diploid level (Jansky et al., 2016; Lindhout et al., 2011). This can be routinely achieved through pollination of a tetraploid of interest with select clones that act as a Haploid Inducer (HI). In a 4x by 2x HI cross, a fraction of the progeny are 2n=2x=24 primary dihaploids lacking chromosomes from the HI parent. Several accessions of diploid Andigenum group potato (formerly *S. phureja* (Spooner et al., 2014)) were demonstrated to act as efficient HIs over half a century ago (Gabert, 1963), and subsequent breeding efforts were successful in obtaining more efficient HIs, as well as incorporating a dominant marker that aids in distinguishing dihaploids from hybrids, which can be triploid or tetraploid, but are typically discarded (Hermsen and Verdenius, 1973; Hutten et al., 1993).

Relatively little is known about the molecular basis of haploid induction in potato, but cytological evidence provides some clues. In a 4x WT by 2x HI cross, dihaploids originate from seeds with hexaploid (6x) endosperm, which is the expected outcome of a 4x central cell fertilization by a 2x sperm (Wangenheim et al., 1960). In different HI clones, 30-40% of pollen fails to complete the second mitosis, resulting in a single, larger restitution sperm, or occasionally, two sperms that may have unbalanced chromosome sets (Montelongo-Escobedo and Rowe, 1969; Montezuma-de-Carvalho, 1967). Furthermore, colchicine-treated pollen, but not untreated pollen of *S. tarijense,* develops restitution sperm and induces potato dihaploids, and colchicine treatment of HI pollen further increases the haploid induction rate (Montelongo-Escobedo and Rowe, 1969). Unreduced 2x sperm are also produced from restitution of the first or second meiotic divisions, but this increased rate of meiotic restitution is not associated with increased haploid induction (Peloquin et al., 1996; Hermsen and Verdenius, 1973). From these results, it was concluded that the 2x sperm fertilizes the central cell, leaving no sperm to fertilize the egg, which then develops parthenogenetically. However, HI-specific DNA markers in dihaploids or near-dihaploid aneuploids were reported in some studies (Clulow et al., 1991; Waugh et al., 1992; Clulow et al., 1993; Wilkinson et al., 1995; Clulow and Rousselle-Bourgeois, 1997; Pham et al., 2019; Bartkiewicz et al., 2018; Ercolano et al., 2004; Straadt and Rasmussen, 2003; Allainguillaume et al., 1997), but not others (Samitsu and Hosaka, 2002; Amundson et al., 2020). This suggests retention of HI DNA either as chromosomes or segments thereof in an otherwise haploid plant, a diagnostic feature of uniparental chromosome elimination in plants (Zhao et al., 2013; Tan et al., 2015; Kuppu et al., 2015; Gernand et al., 2005; Riera-Lizarazu et al., 1996; Laurie and Bennett, 1986; Kynast et al., 2001; Li et al., 2017; Ishii et al., 2010; Maheshwari et al., 2015). In Arabidopsis haploid induction systems, selective instability of the HI chromosomes are observed among the hybrid byproducts (Kuppu et al., 2015; Tan et al., 2015; Maheshwari et al., 2015).

Given the role of dihaploids in efforts to convert potatoes to a diploid inbred crop, the incidental transfer of HI DNA remains a concern, as it has been reported to influence the phenotype of dihaploids (Allainguillaume et al., 1997). The relatively small sample sizes of previous studies, including our previous evaluation of 167 dihaploids that did not identify instances of HI DNA transfer (Amundson et al., 2020), warrant a robust characterization of the frequency and molecular state of incidental HI DNA transfer in primary dihaploids. Furthermore, to our knowledge, no study has investigated the genomic integrity of triploid and tetraploid hybrids obtained from potato haploid induction crosses. Toward these ends, we asked two questions: 1) How often, if ever, do potato HIs transmit chromosomes or chromosome fragments to dihaploids? 2) Do dihaploids or hybrid byproducts of potato haploid induction exhibit evidence of genome instability? In this study, we used genome resequencing to search for HI DNA and determine its molecular state in 1,001 dihaploids and 134 hybrids obtained from potato haploid induction crosses. Among dihaploids, 0.87% of individuals were found to contain HI DNA from 1-3 chromosomes, some of which appear fragmented due to genome instability. Observations of chromosome breakage in dihaploids and hybrids suggest an association between haploidization and genome instability. However, this instability appears ploidy-dependent: HI chromosomes were fragmented much less often in triploid hybrids than in either dihaploids or tetraploid hybrids. Comparison with tetraploid self-pollinated progeny suggested that HI genome instability observed in tetraploid hybrids is not attributable to pollen, sperm or embryo ploidy *per se*. In summary, our results indicate low levels of HI DNA contribution to dihaploids, and suggest a role for ploidy of the HI gamete in potato haploid induction.

## Results

### Widespread aneuploidy in the dihaploid population

To generate dihaploid and hybrids, we pollinated 19 tetraploid clones with haploid inducers IVP35, IVP101 or PL4, recorded presence or absence of inducer-specific anthocyanin pigmentation at the nodes of the plantlets, and evaluated the ploidy of each plantlet by chloroplast counting or flow cytometry (Supplementary Dataset S1). Next, 1,001 individuals from the putatively dihaploid progeny were selected for chromosome dosage analysis by low coverage whole genome sequencing, as previously described (Amundson et al., 2020). Briefly, for each dihaploid, read depth per chromosome is standardized to that dihaploid’s tetraploid parent such that values near 1, 2 or 3 correspond to monosomy, disomy or trisomy, respectively. Aneuploids are then identified as individuals with one or more outlier chromosomes, as shown for a representative cohort of 229 putative dihaploids (Fig. 2A). In this cohort, we identified twenty-seven aneuploids: 25 had one additional chromosome (2n=2x+1=25), one had two additional chromosomes (2n=2x+2=26) and one had three chromosomes (2n=2x+3=27).

**Figure 1.**
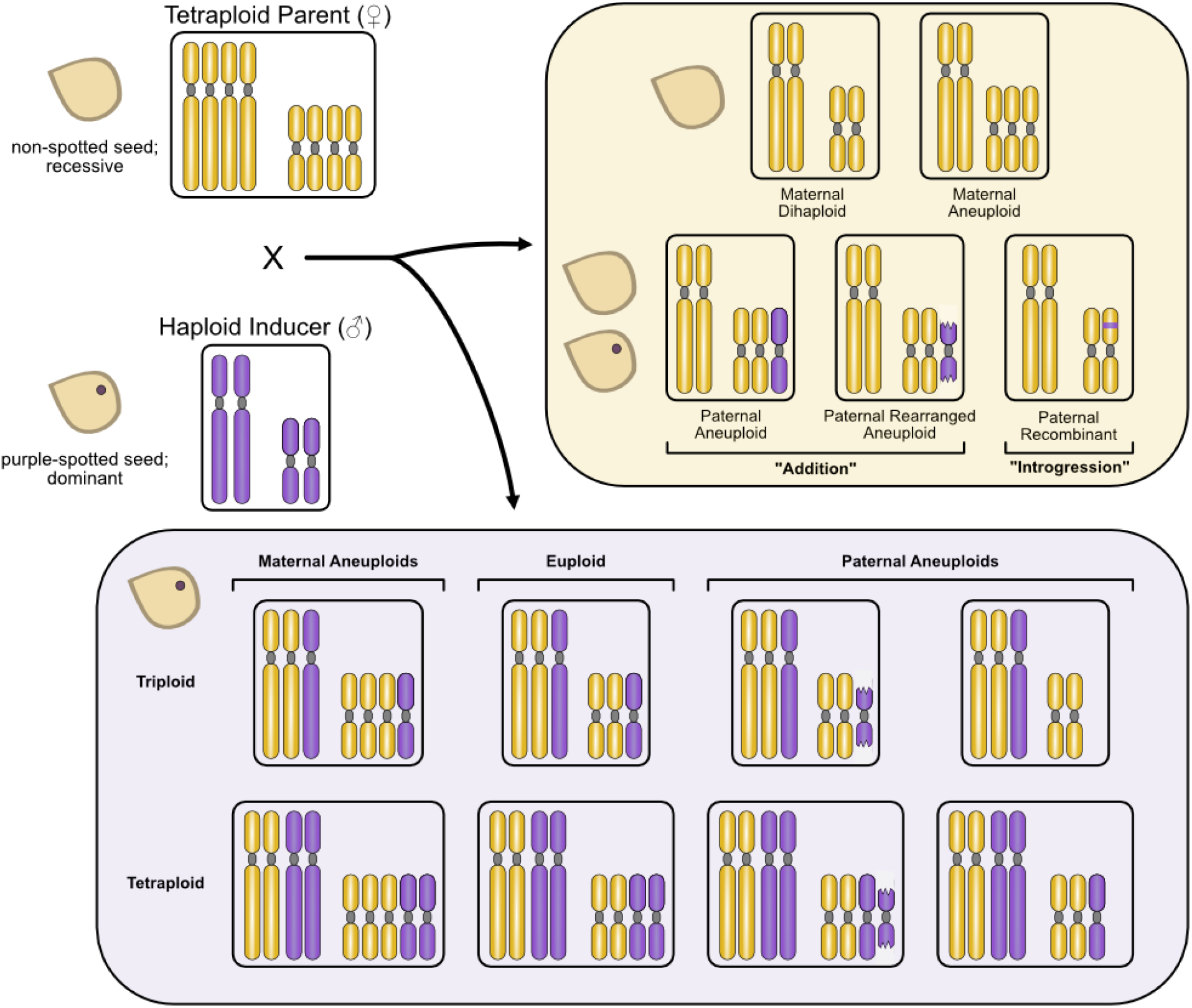
Possible outcomes of potato haploid induction crosses. The haploid-inducing pollinator genotypes used in this study were homozygous for the dominant embryo spot trait. Presence or absence of haploid inducer DNA in the ensuing progeny is expected to manifest as presence or absence of the embryo spot. Progeny from spotted seeds include maternal 2n=2x=24 dihaploids and 2n=2x+1=25 aneuploids. If potato haploid induction is due to post-zygotic elimination of paternal chromosomes, then occasional failure to eliminate all paternal DNA is expected to result in additional paternal chromosomes, intact or rearranged (addition) and/or integration of paternal DNA segments into maternal chromosomes (introgression). Spotted seeds presumably contain the haploid inducer genome and may be triploid or tetraploid depending on the ploidy of the sperm that took part in fertilization. Diploid sperm may result from unreduced pollen or blockage of generative cell division (Montelongo-Escobedo and Rowe, 1969).

**Figure 2.**
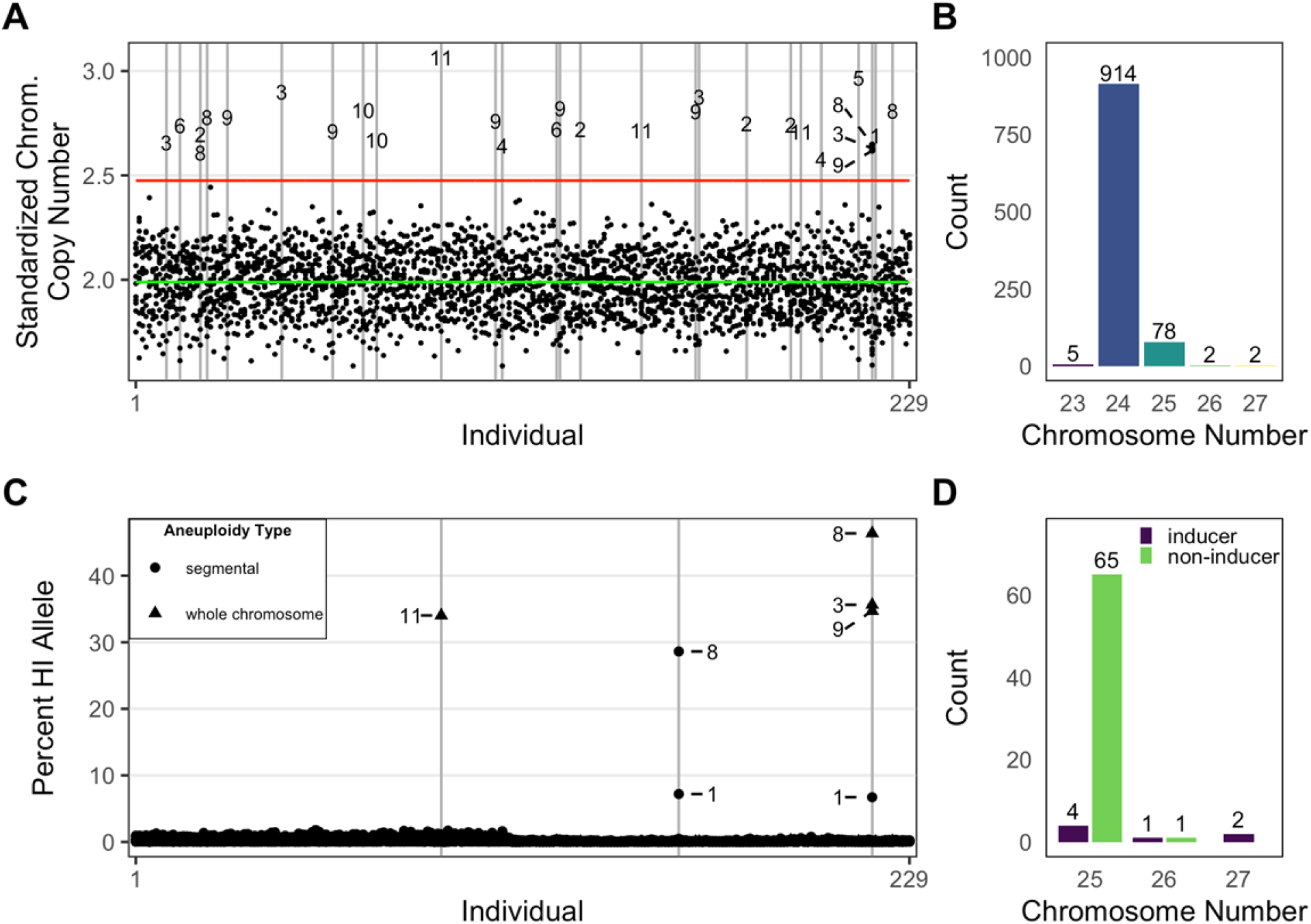
Incidence and parental origin of aneuploidy among putative dihaploids. **A)** Standardized chromosome coverage of 229 dihaploids (inferred from flow cytometry) that were extracted from CIP315047 (WA.077). Each individual is displayed along the X-axis, with the stack of 12 points at each coordinate along the X-axis corresponding to the estimated copy number of the 12 potato chromosomes. The green line corresponds to the population all-chromosome mean, and the red line a cutoff of 3 standard deviations greater than the mean, which was our criterion for calling whole-chromosome aneuploidy. Outliers in this distribution correspond to additional chromosomes, all of which are numbered by homolog. **B)** Count of dihaploids by chromosome number inferred from low pass sequencing for all dihaploids evaluated for chromosome dosage (n=1,001) in this study. **C)** Per-chromosome haploid inducer allele contribution of the 229 flow-cytometry confirmed dihaploids shown in panel A. Each individual is displayed along the X axis as a stack of 12 points, with each point corresponding to the haploid inducer allele contribution of one of the 12 chromosomes. Outliers are numbered by chromosome. Chromosomes identified as outliers in panel A are labeled as whole-chromosome aneuploids; those not identified as outliers are labeled as segmental aneuploids. **D)** Parental origin of chromosomal deficiencies and excesses for all near-dihaploid aneuploids analyzed for parental origin in this study (n=73). All compound trisomics resulted in inheritance of multiple additional chromosomes from the same parent, i.e., a 26-chromosome individual exhibited additional chromosomes from either the maternal or paternal parent, but not both.

Overall, among the 1,001 sequenced putative dihaploids, 8.7% were aneuploid. Primary trisomics (i.e., single chromosome aneuploids) composed 89% of the aneuploid class, with the remaining aneuploids consisting of monosomics (2n=2x-1=23) and primary trisomics for multiple chromosomes (2n=2x+2=26 or 2n=2x+3=27) (Fig. 2B). Each of the 12 homologous chromosomes was recovered as a trisomic. Neither trisomy (chromosome gains only) nor aneuploidy in general (gains and losses pooled) showed significant bias for a particular chromosome (gains only *p*=0.07652, df=11; gains and losses pooled *p*=0.1354, df=11) (Supplementary Fig. S1). It is worth noting that flow cytometric analyses did not readily detect aneuploidy in primary dihaploids.

To evaluate the effect of parental genotype on aneuploidy frequency, we grouped dihaploids based on the genotypes of the parents. When grouped by maternal genotype, aneuploidy frequency ranged from 6.4% to 11.3% and differences between maternal genotypes were not significant (*p*=0.5987; df=6) (Supplementary Fig. S2). When grouped by paternal genotype, aneuploidy frequency ranged from 7.6% to 10.3% and differences between inducer genotypes were not significant either (*p*=0.3626; df=2) (Supplementary Fig. S3). Taken together, and consistent with previous reports (Amundson et al., 2020; Pham et al., 2019; Samitsu and Hosaka, 2002; Wagenvoort and Lange, 1975), our data show that approximately 8.7% of presumed dihaploids are aneuploid, without detectable aneuploidy bias for parental genotype or homologous chromosome in this material.

### Retention of haploid inducer chromosomes

To determine the parental origin of the additional chromosomes in the aneuploid dihaploid progeny, we identified homozygous SNPs between each pair of tetraploid seed parent and haploid inducer. We used these SNPs to calculate the percentage of reads that originated from the haploid inducer across the genome of every dihaploid. If the additional chromosome originated from the haploid inducer, this percentage is expected to be approximately 33% while it will remain close to 0% if all copies originated from the tetraploid parent. Representative SNP dosage plots are shown in Fig. 2C. In this population, two of the aneuploids identified in Fig. 2A carried chromosomes from the haploid inducer parent. One of these two individuals also exhibiting haploid inducer alleles above background levels on chromosome 1 (Fig. 2C). A third individual was not aneuploid according to dosage analysis but showed haploid inducer alleles above background levels on chromosomes 1 and 8 (Fig. 2C).

In total, we analyzed SNP dosage for the 923 dihaploids for which we had sufficient SNP information. Of those, 74 had extra chromosomes or large chromosomal segments, eight of which were derived from the HI; all others were of maternal origin (Fig. 2D). Two lines, MM247 and MM890, were segmental aneuploids with additional HI chromosome segments; the six others showed HI-derived aneuploidy of entire chromosomes (Fig. 3; Supplementary Fig. S4). We refer to these eight lines as HI addition dihaploids hereafter. All of the haploid inducer genotypes contributed genetic material to at least one dihaploid, and the frequencies at which they did so were not significantly different (*p*=0.7747; Fisher Exact test). The low overall frequency (8/923; 0.87%) of HI chromosome retention is consistent with previous results in which chromosomes from the inducer parent were not detected in cohorts of less than 200 individuals (Amundson et al., 2020; Samitsu and Hosaka, 2002; Pham et al., 2019).

**Figure 3.**
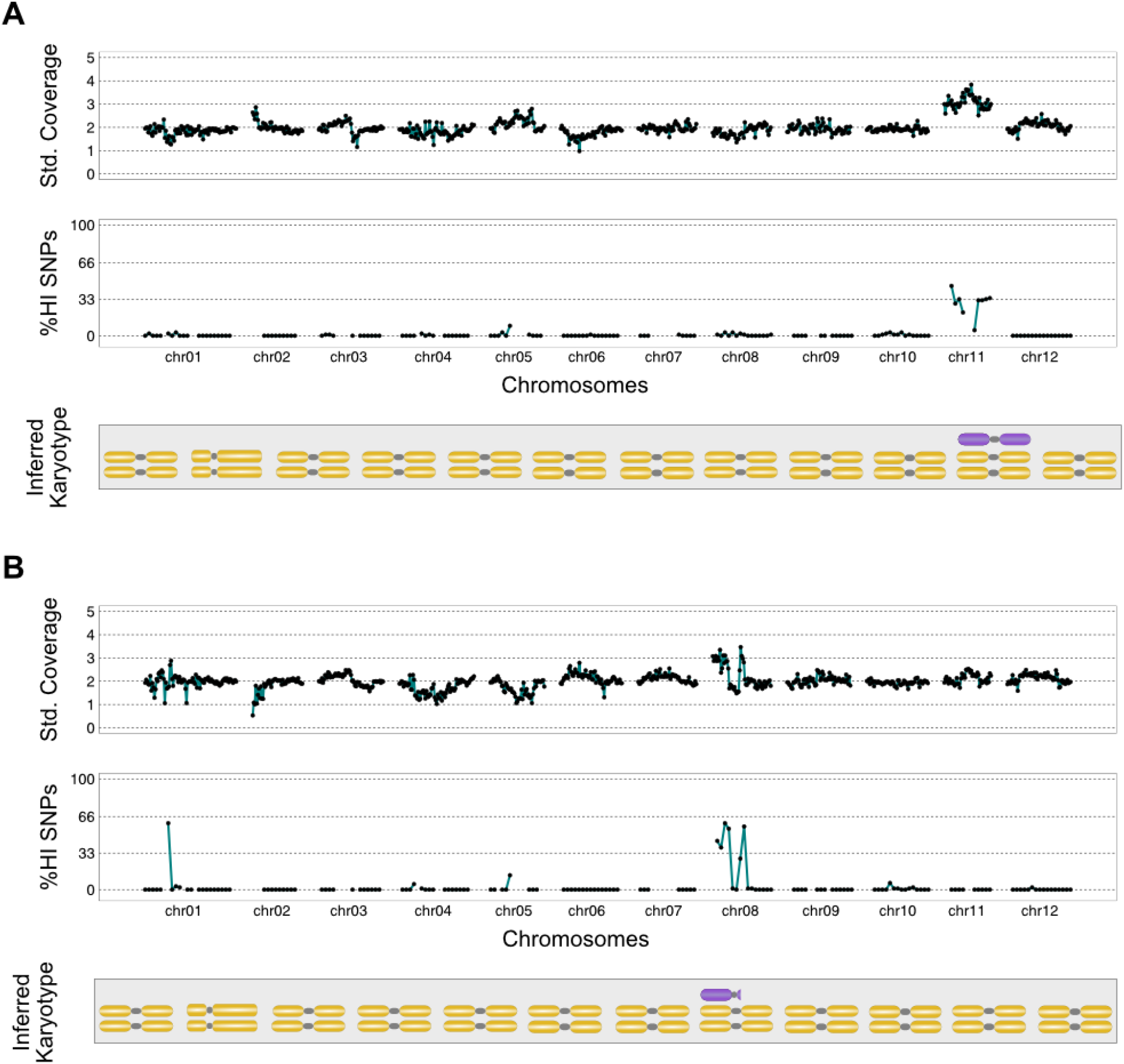
Paternal genomic contributions to maternal dihaploids. *In silico* karyotypes of trisomic dihaploids demonstrate the presence of haploid inducer DNA. **A)** Dihaploid MM246, with additional chromosome 11 from haploid inducer IVP35. **B)** Dihaploid MM247, with segmental aneuploidy of chromosome 8 from haploid inducer IVP35. The HI segmental addition on chromosome 1 cosegregates with chromosome 8 in each of seven dihaploid populations we evaluated in this study, suggesting that the two loci are physically linked and that potato is either polymorphic for Chr8-1 translocation, or that this is an assembly error in the reference genome.

### Detection of haploid inducer-derived DNA segments in dihaploids

Appearance of inducer DNA fragments shorter than entire chromosomes have also been reported among potato dihaploids (Wilkinson et al., 1995; Pham et al., 2019). Our low coverage sequencing cannot detect segments of this size, but they may be detected with higher coverage. To test whether this type of transfer occurred in our material, we sequenced all HI addition lines to higher coverage (depth=19-102x) and surveyed three of them for regions of HI alleles on the disomic chromosomes. Our initial search revealed that, in each dihaploid, the fraction of loci with HI alleles was low (0.39-0.69%) and within the range of previous reports (Pham et al., 2019; Bartkiewicz et al., 2018; Amundson et al., 2020). However, at these loci where HI alleles were detected, the HI alleles were systematically underrepresented in read counts, suggesting that they were artifacts, possibly resulting from sequencing or mapping errors (Supplementary Fig. S5) (Pham et al., 2019; Amundson et al., 2020). We next applied stronger filter thresholds to the entire dataset. After this more stringent filtering, 3-13 tracts of HI alleles remained in each of the HI addition dihaploids (Table 1). Notably, identical tracts were detected in lines MM247 and MM1114, both of which carry part or all of chromosome 8 from IVP35. One of these tracts, a 1.8 Mb region of chromosome 1 corresponds to a scaffold that appeared linked to chromosome 8 in three independent potato mapping populations (Endelman and Jansky, 2016; Bourke et al., 2015) as well as in seven of our dihaploid populations (Supplementary Fig. S6) suggesting pre-existing translocation in the genome of DM1-3 or an error in the version of the DM1-3 genome assembly that was used here (version 4.04). In conclusion, our analysis does not provide evidence of true introgressions of short HI DNA segments.

**Table 1:** Putative introgressions of haploid inducer (HI) DNA segments in dihaploid potatoes. For each HI addition dihaploid, the trisomic chromosome and reference genome coordinates of putative segmental introgressions are shown.

| HI Addition Dihaploid | Tetraploid Parent | HI Parent | Addition Chromosome(s) | Segments of HI alleles |
| --- | --- | --- | --- | --- |
| MM246 | WA.077 | IVP35 | Chr11 | Chr01:45392940-45400039 |
|  |  |  |  | Chr05:15510198-15510533 |
|  |  |  |  | Chr07:7833610-7844546 |
|  |  |  |  | Chr07:7880027-7880413 |
| MM247 | WA.077 | IVP35 | Chr08 | Chr01:24228918-25308116* |
|  |  |  |  | Chr01:25309455-26003606 |
|  |  |  |  | Chr01:38172625-38174004* |
|  |  |  |  | Chr07:9467446-9467874 |
| MM1114 | WA.077 | IVP101 | Chr03, Chr08, Chr09 | Chr01:24228918-26003606* |
|  |  |  |  | Chr01:38172625-38174004* |
|  |  |  |  | Chr01:41494766-41499334 |
|  |  |  |  | Chr01:67944376-67945914 |
|  |  |  |  | Chr01:84519844-84521775 |
|  |  |  |  | Chr05:280661-281197 |
|  |  |  |  | Chr05:901127-901352 |
|  |  |  |  | Chr05:15812350-15814025 |
|  |  |  |  | Chr07:5953200-5953784 |
|  |  |  |  | Chr07:9467446-9467874* |
|  |  |  |  | Chr10:52856295-52859220 |
|  |  |  |  | Chr10:54984736-54988488 |
|  |  |  |  | Chr12:55689534-55690262 |

\* Coordinates of putative introgression common to multiple individuals.

### Selective instability of the haploid inducer genome in dihaploids and tetraploid hybrids

Next, we asked whether the hybrid byproducts of potato haploid induction showed signs of genome instability. Seeds with the dominant, inducer-specific embryo spot marker were germinated and analyzed by flow cytometry, yielding 30 triploids and 104 tetraploids. As a control, we included 14 progeny that did not show the nodal banding phenotype, were tetraploid by flow cytometry, and lacked HI alleles in the low coverage sequencing; these are likely self-pollinated progeny (selfs) of the tetraploid clones. To distinguish novel dosage variants attributable to genome instability from recurring variants likely due to pre-existing structural variation in the parents, each offspring was evaluated in the context of its siblings of the same ploidy (Fig. 4A-B). Aneuploids made up a greater proportion of tetraploid hybrids (>70% vs 22% of triploid hybrids), with the frequency of maternally and paternally derived aneuploidy both increasing (Fig. 4C). The per-chromosome rate of HI-derived segmental aneuploidy was significantly lower in the triploids hybrids than the corresponding rate in either dihaploids or tetraploid hybrids, suggesting a greater degree of HI genome instability in dihaploids and tetraploid hybrids (Table 2). Chromosome breakage was not observed in tetraploid selfs, suggesting that the instability of HI-derived chromosomes seen in tetraploid hybrids was not a consequence of 2x pollen and/or sperm *per se* (Table 2). Relative to triploid hybrids, tetraploid hybrids showed a strong and highly significant increase in the incidence of HI-derived genome instability (*p*<0.001; log odds 95% CI 2.62247-1.501144) (Fig. 4C), a difference driven by more frequent segmental aneuploidy of HI-derived chromosomes (Fig. 4D). Most tetraploid hybrids exhibited no more than two novel CNV of each parental genome, indicating that genome instability is not restricted to few exceptional tetraploid hybrids, but is pervasive (Fig. 4E). In conclusion, our data suggest that HI-derived chromosomes are selectively unstable in dihaploids and tetraploids, suggesting a specific role of 2x HI sperm in potato haploid induction.

**Table 2:**
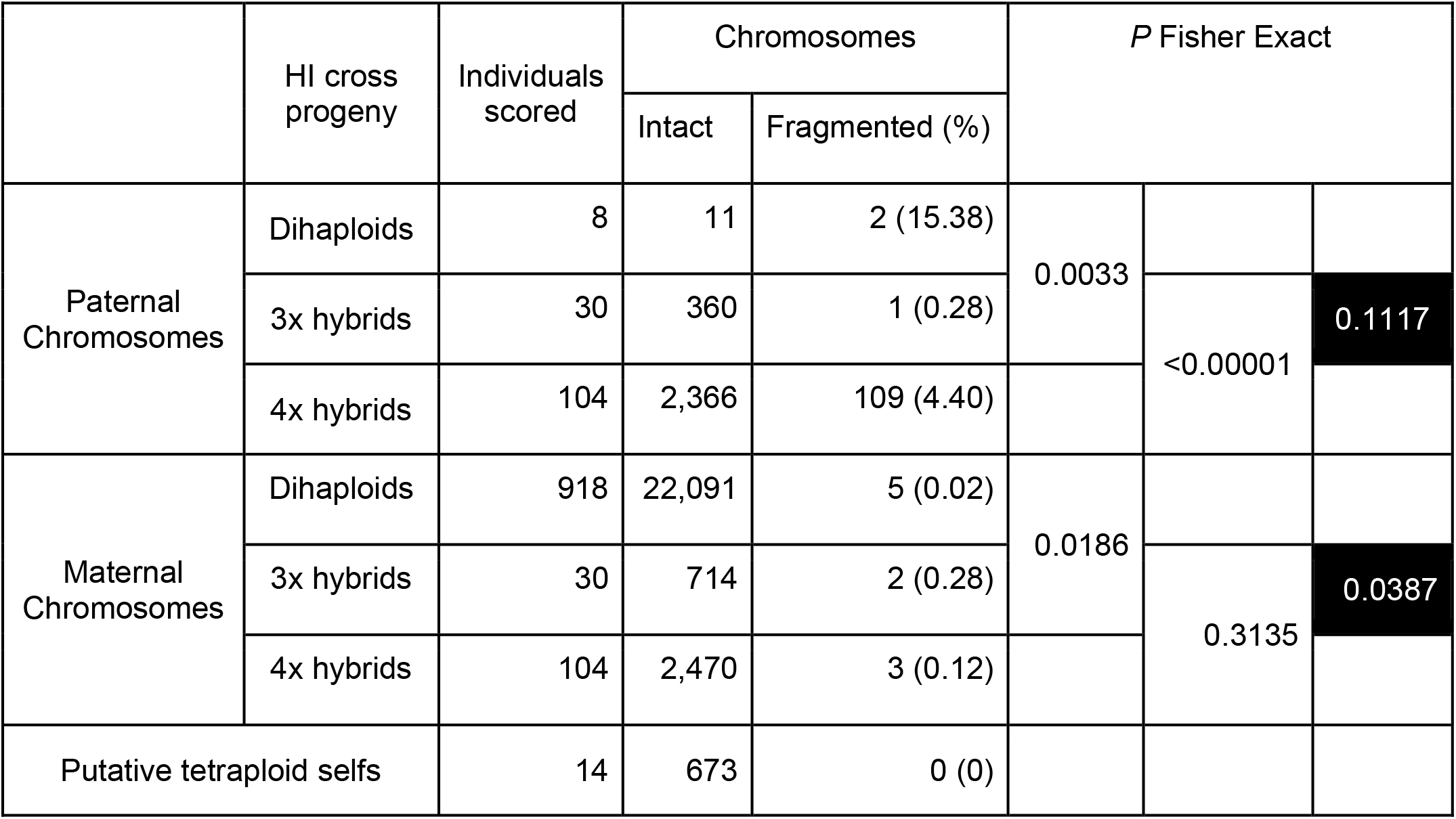
Frequency of chromosome breakage among progeny of potato haploid induction crosses. For each progeny class, maternal and paternal chromosomes were recorded as appearing in an intact or fragmented state. The number and frequency of chromosomes of each type (intact vs. fragmented) were then grouped by rogeny ploidy and parental origin.

**Figure 4.**
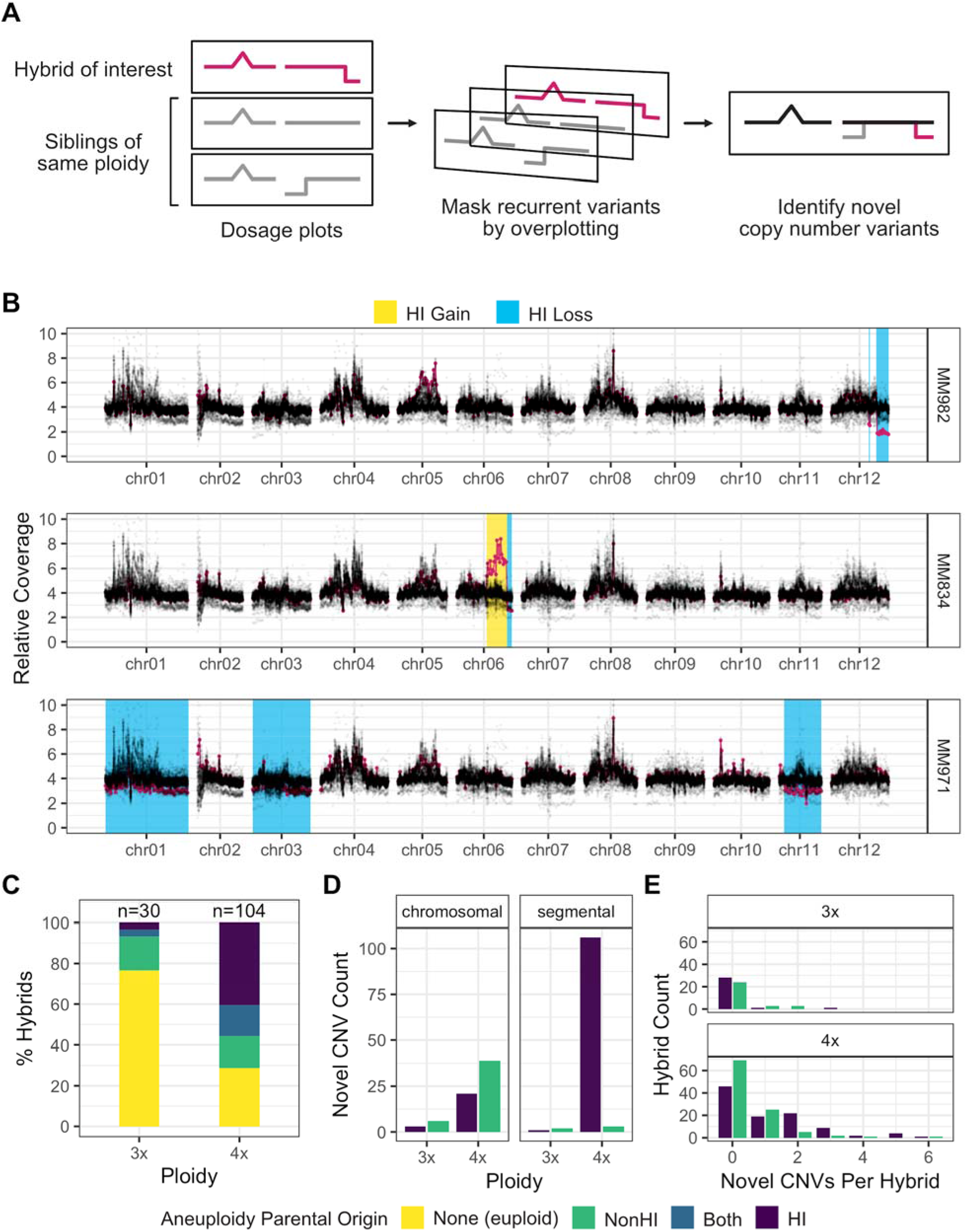
Haploid inducer (HI) genome instability in tetraploid potato hybrids. Chromosomal variation was investigated by dosage analysis in triploid and tetraploid hybrids. **A)** Schematic of dosage plot generation for a hybrid cohort. An individual displaying novel dosage variation is identified as an outlier track (pink) compared to common structural variation (gray) in overlaying dosage plots obtained by plotting siblings of the same ploidy. Parental origin of each novel variant is inferred from allele-specific read depth at parent-informative SNP loci. **B)** Overlay plots of the same sibling hybrid family with novel copy number variations (CNV) highlighted. Regions corresponding to DNA gains or losses are shaded with yellow or blue backgrounds, respectively. In these examples, all novel CNV are attributable to gained or lost haploid inducer DNA. **C)** Increased paternal aneuploidy in near-tetraploid vs. near-triploid offspring. Combined data from all cohorts in this study. **D)** Paternally derived segmental variation is preponderant in tetraploid hybrids. The bars display counts of aneuploidy according to paternal origin and ploidy of individuals. They also display the counts of whole chromosome aneuploidy vs segmental aneuploidy. As only haploid inducer chromosome breaks are considered in this panel, both classes may exhibit whole chromosome aneuploidy of either parent and segmental aneuploidy of non-inducer chromosomes in addition to HI chromosome breakage. **E)** Number of novel CNV events per hybrid, subdivided by parental origin, showing that haploid inducer genome instability in hybrids is not restricted to few individuals.

### Tetraploid hybrids produced by first meiotic division restitution of the haploid inducer

As restitution sperm formation, but not 2n gamete, appears to be associated with potato haploid induction (Montelongo-Escobedo and Rowe, 1969; Peloquin et al., 1996; Dongyu et al., 1995), we asked which type of 2N gamete gave rise to our tetraploid hybrids and which, if any, was associated with HI genome instability. From the high-coverage sequencing of each dihaploid addition line, we derived the HI haplotypes of the additional chromosome(s) in each line (Fig. 5A), which we refer to as H’ hereafter. Using these haplotypes, we then used the low-pass sequencing of 134 hybrids to genotype the centromeres of the chromosomes contributed by the HI. As a control, we analyzed triploid hybrids and found the expected transmission of a single HI haplotype through the centromere and into the chromosome arms (Supplementary Fig. S7), indicating that our centromeric HI haplotype phasing was robust. For tetraploid hybrids, the HI-contributed sequences at centromere-linked markers are expected to be heterozygous if derived from 2n first division restitution (FDR) pollen, but homozygous if derived from either 2n second division restitution (SDR) pollen or 2x restitution sperm (Fig. 5B). Among 78 tetraploid hybrids, all but five showed HI heterozygosity at the centromeres, implicating FDR as the dominant mechanism of hybrid formation (Fig. 5C; Supplementary Fig. S8). Among the five hybrids with the SDR or RS pattern, one showed ~50% H’ allele of Cen11, but this individual was disomic for chromosome 11 missing both maternal homologs (Supplementary Fig. S9); together, these results are also consistent with FDR. Of the remaining four hybrids, all of which were derived from IVP101, two showed signs of HI genome instability and two did not. In conclusion, while FDR hybrids predominated among tetraploids, with minor contributions possibly from SDR gametes or restitution sperm, no mechanism was uniquely associated with HI genome instability.

**Figure 5.**
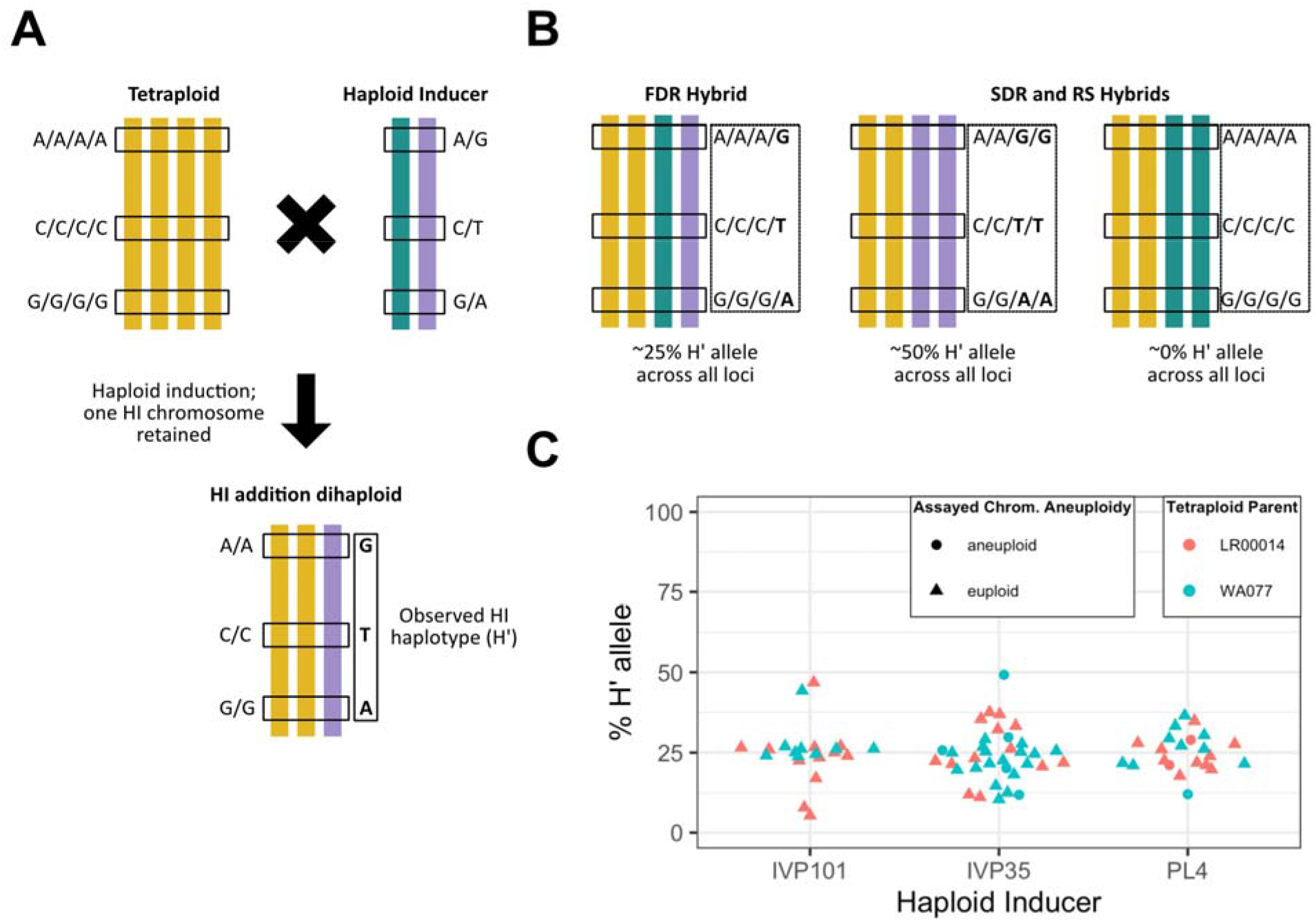
Tetraploid hybrid formation in potato haploid induction. **A)** Biallelic SNP loci used for analysis are homozygous in the tetraploid parent and heterozygous but unphased in the HI. The retained HI chromosome(s) in each HI addition dihaploid represents a phased HI-derived haplotype. To avoid confounding effects from crossovers, only expectations and data for the non-recombining centromere of is shown. **B)** Expected representation of HI haplotype alleles in a non-recombining region of tetraploid hybrids. For all tetraploid hybrids, read information adjacent loci is binned and the percentage of reads with H’ alleles is calculated. If tetraploids are the product of first division restitution (FDR) of the HI, then H’ alleles are expected to appear in 25% of all binned reads. If tetraploids are the product of second division restitution (SDR) or restitution sperm (RS), the expected percentage could be 50% or 0% depending on which HI haplotype was inherited. **C)** Percentage of H’ allele in the centromeres of tetraploid hybrids. Each point corresponds to the percentage of H’ allele among reads spanning the non-recombining region of a chromosome of one tetraploid hybrid. Chromosome 8 was used to assess IVP101 hybrids. Chromosome 10 was used to assess PL4 hybrids. Chromosome 11 was used to assess IVP35 hybrids.

## Discussion

Multiple studies investigating the presence of haploid inducer genetic material in potato dihaploids have come to different conclusions (Amundson et al., 2020; Pham et al., 2019; Bartkiewicz et al., 2018; Ercolano et al., 2004; Straadt and Rasmussen, 2003; Samitsu and Hosaka, 2002; Allainguillaume et al., 1997; Clulow and Rousselle-Bourgeois, 1997; Wilkinson et al., 1995; Clulow et al., 1993; Waugh et al., 1992; Clulow et al., 1991). The extent of HI DNA contamination is important both for a basic understanding of haploid induction and for applied utilization of primary dihaploids for diploid potato breeding. We previously argued that, based on characterized haploid induction systems, whole chromosomes or large fragments should be the most likely form of inherited HI DNA (Amundson et al., 2020). Smaller introgressions could also occur (Pham et al., 2019), but they need robust validation to avoid the confounding effect of polymorphism in the tetraploid seed parent and pervasive sequencing or alignment artifacts.

Previously, we found that 167 dihaploids were free of haploid inducer DNA. It is possible, however, that low-frequency HI-DNA transmission was missed. To address this possibility, we scaled up our sampling and obtained sequencing of 1,001 putative dihaploids. Based on the availability of robust polymorphic markers, we identified HI DNA in 8 individuals out of 923, 6 with whole chromosomes and 2 with large fragments. The presence of HI chromosomes or fragments in 0.87% of dihaploids is good news for potato breeders because they can expect most of their haploids to be devoid of HI DNA.

We documented the occasional appearance of small HI segments (0.5 to few kb). The evidence in support of their presence is robust because it is based on a continuous haplotype that encompasses multiple SNPs. These could represent small translocations or gene conversion derived from the HI genome before elimination. However, identical HI segments were present in HI addition lines MM247 and MM1114, which were both trisomic for chromosome 8. Part of chromosome 8 was previously identified as translocated or misassembled in at least two genetic mapping populations (Endelman and Jansky, 2016; Bourke et al., 2015) as well as in our dihaploid populations. Therefore, a simpler interpretation is that these short introgressions are actually part of the addition chromosome forming the associated trisomy. They appear as isolated regions due to genome assembly artifacts or pre-existing translocations in the HI relative to the reference genome. This was demonstrated when two independent additions of the same chromosome were available, and seems the best explanation for all instances. Further study, including local assembly of translocations or conversion segments with flanking DNA will be needed to clarify the nature of these events.

The presence of HI chromosomes in some dihaploids could be explained if, after formation of a hybrid genome, the HI genome was imperfectly eliminated. Acceptance of this hypothesis would imply that uniparental genome elimination is the mechanism that generates haploids and that the dihaploids with no HI DNA contamination underwent perfect elimination of the HI genome.

There are, however, at least two alternative explanations. First, that all or a fraction of the perfect dihaploids result from parthenogenesis. This would require two independent mechanisms of haploid induction to work in the potato system, and seems implausible. Second, that the mechanism of haploid induction entails incomplete gamete fusion. A defective sperm may thus deliver both egg-activating factors and, occasionally, chromosomes, but fail to carry out proper karyogamy. This mechanism has not been explicitly proposed, to our knowledge, but may explain observations with the maize haploid induction system, where in addition to phospholipase, mutation of fusogenic proteins both enhance HI rate, and induce haploids in the wild-type phospholipase background (Zhong et al., 2020, 2019).

What fates are possible for the HI genome? The genome delivered by sperms formed by the tetraploid selfs were stable. We could assess HI genome integrity in HI-contaminated dihaploids, and in triploid and tetraploid hybrids. Dihaploids that had inherited and maintained HI chromosomes shared high instability of the HI genome with tetraploid hybrids. Triploids, on the other hand, did not. This demonstrates that the HI genome can be inherited and maintained with fidelity and that instability is not intrinsic to the formation of hybrid zygotes. The tetraploids result from hybridization of a 1n(=2x) egg sac with 2n(=2x) sperm. We infer that either during formation of the 2n sperm or upon fertilization, the HI genome becomes unstable. The instability displayed by tetraploid hybrids could be related to that displayed by HI-containing dihaploids. This provides a potential explanation for the long-standing proposal that 2n sperm triggers haploid induction (Montelongo-Escobedo and Rowe, 1969; Montezuma-de-Carvalho, 1967; Wangenheim et al., 1960). These studies documented the formation of 2x sperm from restitution of the generative cell mitosis in pollen of HIs and suggested a connection to haploid induction. Genome maintenance may become compromised during formation or growth of 2n pollen resulting in a fragmented genome that is incompetent for replication and subject to elimination.

It is also possible that instability is unrelated to genome elimination. We explored the nature of 2n sperms in the HI crosses by analyzing the HI contribution in tetraploid hybrids. Restitution of second mitosis predicts perfect homozygosity of 2n sperm. Instead we found heterozygosity indicative of meiotic First Division Restitution (FDR) in most cases. This leaves unanswered the role of 2x sperm in HI. It demonstrates, however, that the instability observed in the tetraploid hybrids is connected to FDR. This instability could result from missegregation during meiosis (Umbreit et al., 2020), but its relation to HI is mysterious.

In conclusion, using a large-scale approach, we examined the genome of 1135 progeny from HI crosses determining that, regardless of parental genotype, a small but definite fraction of dihaploids display paternal HI contribution. This large-scale study provides the solid evidence needed to interpret previous studies, and calibrates the expectations for potato HI crosses. In addition, we made an unexpected observation. The HI genome, which is stable when inherited by triploid hybrids, displays selective instability both in tetraploid hybrids and in dihaploids. The interpretation and meaning of these findings are still open. At a minimum, they indicate that genome stability is compromised in the HI 2n pollen. They also reinforce the hypothesis that 2n pollen may be required for HI, suggesting future lines of investigation to elucidate mechanisms contributing to this unusual, but highly relevant phenomenon.

## Materials and Methods

### Plant material

Primary dihaploids and hybrids were obtained from 19 tetraploid clones (Supplementary Table S2) *via* pollination with haploid inducers IVP101 (Hutten et al., 1993), IVP35 (Hermsen and Verdenius, 1973) or PL4 (also known as CIP596131.4; Ordoñez et al, in prep) in greenhouses located at the CIP’s experimental station in the Peruvian Andes (3,216 masl, −12.01039, - 75.22411). Flower buds of the pistillate parents were emasculated and pollinated with HI pollen the following day. All haploid inducers are homozygous for a dominant embryo spot that facilitates the detection of hybrids (Hermsen and Verdenius, 1973; Hutten et al., 1993). Seeds were extracted from mature fruit, recorded for presence or absence of the embryo spot, and germinated on soil. The ploidy of each established seedling was determined by either chloroplast counting as described in (Amundson et al., 2020) or flow cytometric measurement of nuclear DNA content against maternal and paternal parents as standards. Refer to Supplementary Dataset 1 for an expanded description of plant material.

### Flow cytometry

Approximately 50-60 mg of greenhouse-grown leaf tissue was harvested from each sample and homogenized in 500 μl of LB01 buffer (Doležel et al., 1989) and left to rest for 1 minute. 250 μl of homogenate was passed through a 20μm filter (Partec 04-0042-2315) into tubes containing 12 μl of 1mg/ml propidium iodide and 2.5μl of 5mg/ml RNase. Samples were incubated in the dark for 5 minutes and analyzed in an Accuri C6 flow cytometer (BD biosciences) with the following filter configurations: a) FL-1 530/14-nm bandpass filter, b) FL-2 585-20nm bandpass filter and c) FL-3 670-nm longpass filter. Threshold levels were set to 10,000 for forward scatter (FSC) with a secondary threshold of 1,000 for FL-2 (Galbraith et al., 2011).

### Whole genome resequencing

Genomic DNA was extracted from young leaflets as previously described (Ghislain M., Zhang D. P., Herrera, M. R., 1999). For each sample, approximately 750ng of genomic DNA was sheared to an average size of 300bp as previously described (Amundson et al., 2020). Libraries were constructed using all sheared input DNA with KAPA Hyper Prep kit (cat. No KK8504) with half-scale reactions used throughout the protocol, custom 8bp index adapters, and amplification cycles as described in Supplementary Dataset 2. Libraries were sequenced on Illumina HiSeq 4000 or NovaSeq 6000 platforms at the University of California, Davis DNA Technologies Core, Vincent Coates Genome Sequencing Laboratory, or University of California San Francisco Center for Advanced Technologies, as specified in Supplementary Dataset 2. Libraries were demultiplexed using custom Python scripts available on our laboratory website (allprep-12.py; http://comailab.genomecenter.ucdavis.edu/index.php/Barcoded_data_preparation_tools). Publicly available sequencing reads from (Pham et al., 2017), (Hardigan et al., 2017) and (Amundson et al., 2020) were retrieved from NCBI Sequence Read Archive and incorporated in subsequent analyses.

### Variant calling

Adapter and low quality sequences were trimmed from raw reads using Cutadapt v1.15 (Martin, 2011), retaining reads ≥ 40nt in length. Trimmed reads were aligned to the DM1-3 v4.04 reference assembly, including DM1-3 chloroplast and mitochondrion sequences, using BWA mem (v0.7.12r1039) and default settings (Li, 2013). Alignments were further processed to remove PCR duplicates, soft clip one mate of overlapping read pairs, remove read pairs with mates aligning to different chromosomes, and locally realign indels, as previously described (Amundson et al., 2020). Processed alignments were then used as input for joint variant calling and genotyping with FreeBayes (version 1.3.2) (Garrison and Marth, 2012) with minimum mapping quality 20, base quality 20, Hardy-Weinberg priors off, and up to 4 alleles considered per variant, and all other parameters left at the default setting.

Initially, we genotyped a subset of parental clones with deep whole genome sequencing available from this study or from previous studies (Hardigan et al., 2017; Pham et al., 2017), which we designated “Cohort A” in Supplementary Dataset 2. Raw variants were filtered as follows: NUMALT == 1, CIGAR == 1X, QUAL ≥ 20, MQM ≥ 50, MQMR ≥ 50, |MQM - MQMR| < 10, RPPR ≤ 20, RPP ≤ 20, EPP ≤ 20, EPPR ≤ 20, SAP ≤ 20, SRP ≤ 20. For each pair of tetraploid parent and haploid inducer represented in the offspring of Cohort A, we identified loci with read depth within 1.5 times the genome-wide median of each parent, at least 10 supporting reads, and homozygous genotype calls for different alleles in the two parents. Each list of parental SNPs was used to provisionally determine chromosome dosage (see ‘Chromosome Dosage Analysis’ below) for all offspring of Cohort A. To determine parental origin of aneuploidy in offspring for which sequencing reads from the tetraploid parent were not available, we tested the possibility of using pooled reads from multiple dihaploids produced from the same parent instead. Specifically, we tested the effect of substituting pooled low-coverage sequencing from dihaploids for that same tetraploid parent at the SNP calling step, and tested if we could recapitulate the observations obtained using SNPs taken directly from the tetraploid parent. As a proof of concept, we pooled low-coverage alignments from 205 dihaploids of WA.077 at the variant calling step and repeated all downstream analysis of Cohort A samples. Upon obtaining acceptable results, pooled alignments from low coverage dihaploids extracted from C93.154 (n=237), 93.003 (n=73), C91.640 (n=79), LR00.014 (n=110), LR00.022 (n=51), LR00.026 (n=51), WA.077 (n=205), and all deeply sequenced HI addition lines were included along with Cohort A samples for the variant calling and genotyping reported in the manuscript.

### Chromosome dosage analysis

Read alignments from low coverage dihaploids and hybrids were filtered for mapping quality ≥10 and counted in non-overlapping 1Mb bins using bedtools (version 2.27.1). We calculated the fraction of all aligned reads that mapped to a chromosome, normalized this fraction to the corresponding fraction of a family-specific control sample (controls specified in Supplementary Dataset 1) and scaled the standardized coverage values to the expected ploidy state based on flow cytometry results. Putative aneuploids were identified as outliers with a standardized coverage value of ≥3 standard deviations from the within-family all-chromosome mean. In some families, segregation of pre-existing deletions on chromosome 12 resulted in a high rate of false positive trisomy and monosomy calls. False positives of this nature are listed in Supplementary Dataset 1. Individuals exhibiting a false positive signal on chromosome 12 were not recorded as aneuploid, unless they also exhibited aneuploidy of another chromosome type. To infer parental origin of numerical and structural aneuploidies, parental SNPs were identified for each combination of tetraploid parent and haploid inducer as described above. For each low-coverage dihaploid or hybrid, allele-specific read depth was then tallied at homozygous parent-informative SNP loci in non-overlapping 4Mb bins, and bins with fewer than 30 reads covering all informative loci within a bin were withheld from analysis.

### High resolution analysis of parental DNA contribution

For each HI addition line, genotype data were recorded as T=tetraploid parent H=haploid inducer. For high stringency filtering, loci were removed from consideration if any of the following criteria were met: i) one or more reads matched the HI allele in the tetraploid parent, ii) three or more reads matched the HI allele in the dihaploid pool, iii) excessive read depth (greater than the mean depth plus four standard deviations greater than the mean depth) was observed in either parent or the dihaploid at hand (Li, 2014), iv) HI allele depth was < 6 in the dihaploid at hand or v) the HI allele represented <15% of the total reads at a locus in the dihaploid at hand.

### Low-pass haplotype analysis

Biallelic SNP loci with homozygous genotype calls for either allele in WA.077, heterozygous genotype calls in IVP35 and heterozygous with a single dose of the HI-specific allele (i.e., 0/0/1 if the called tetraploid genotype was 0/0/0/0 and 0/1/1 if the called genotype was 1/1/1/1) were used to define phased alleles of H’. For each tetraploid hybrid, we then calculated the depth of reads with H’ and non-H’ alleles at all retained loci, aggregated counts across the DM1-3 coordinates defined as recombination-suppressed centromeres by Bourke et al. (2015) or by non-overlapping 4Mb bins, and reported the ratio of reads matching H’ reads to H’ + non-H’ reads.

### Statistical analyses

#### Proportion of aneuploids among dihaploids by female parent

Euploidy and aneuploidy were treated as discrete outcomes, and counts of each category were evaluated for statistical significance using the prop.test() function in R version 3.6.2 (R Core Team 2019). Only families with 30 or more dihaploids were included in the analysis.

#### Fisher exact counts of HI dihaploid introgression events by haploid inducer

Appearance of HI-derived chromosomes in an otherwise dihaploid plant was treated as a binary outcome and used to construct a 2×3 contingency table with each HI genotype. Only dihaploids for which we had sufficient SNP information to determine chromosome parental origin were considered. This included dihaploids from the following tetraploid parents: 93.003 (CIP390637.1), Atlantic (CIP800827), C01.020 (CIP301023.15), C91.640 (CIP3888615.22), C93.154 (CIP392820.1), Desiree (CIP800048), LR00.014 (CIP300056.33), LR00.022 (CIP300072.1), LR00.026 (CIP300093.14) and WA.077 (CIP397077.16). This table was used to conduct a Fisher Exact test in R using the function fisher.test().

#### Linkage disequilibrium from dosage variable states

For each dihaploid, standardized coverage values and bin dosage states were derived for non-overlapping 1Mb bins of the reference genome as previously described (Amundson et al., 2020). Fisher Exact tests were then carried out between pairs of dosage states to assess linkage disequilibrium between bins. For example, assume that both Bin1 and Bin100 have three dosage states: standardized coverages 1, 2 and 3. To test whether Bin1-State1 was correlated with Bin100-CN3, the following four dihaploid sets were compared in a 2×2 contingency table: *observed in Bin1-State1: observed not in Bin1-State1, expected in Bin1-State1: expected not in Bin1-State1*, where the expectation was derived from the assumption of complete independence. Self-comparison and reciprocal comparisons were removed, and the remaining comparisons were controlled at false discovery rate (FDR) = 0.05 unless otherwise noted. Chromosomal bins in statistically significant linkage disequilibrium (LD) with one another for any state at that pair of bins were displayed in an LD matrix.

#### Logistic regression model for incidence of paternal (maternal) genome instability in triploid and tetraploid hybrids

For each hybrid, we determined the incidence and parental origin of whole-chromosome and segmental aneuploidy tetraploid hybrids as previously described for potato dihaploids (Amundson et al., 2020). Instability of the paternal genome was treated as a binary outcome and used in a logistic regression model with ploidy, genotype of the HI parent and aneuploidy of maternal chromosomes included as predictor variables. Contribution of the maternal genome to aneuploidy of HI-derived chromosomes was determined from the maternal aneuploidy term in the model. Pairwise differences between ploidy levels were evaluated with Tukey multiple test correction. To evaluate the stability of maternal chromosomes, maternal aneuploidy was used as the binary response variable and paternal aneuploidy was incorporated as a predictor variable. Effects of paternal aneuploidy on maternal aneuploidy, as well as ploidy-dependent effects were evaluated as described above.

## Data Availability

All sequencing data generated in this study is currently being deposited at NCBI Sequence Read Archive under a Project ID that remains to be determined, and will be updated as soon as possible. IVP101 whole genome sequencing was retrieved from NCBI Sequence Read Archive project ID PRJNA408137. Whole genome sequencing of cv. “Atlantic” was retrieved from NCBI Sequence Read Archive project ID PRJNA287438. Code for read preprocessing, variant calling, and chromosome dosage analysis is available on https://github.com/kramundson/MM_manuscript. Supplemental Datasets S1 and S2 are available on Dryad at https://doi.org/10.25338/B8JS8D.

## Acknowledgements

We thank Hannele Lindqvist-Kreuze for guidance and support of the CIP-based research and the CIP’s greenhouse technicians who performed most of the crossings and plant maintenance. This work was supported by the National Science Foundation Plant Genome Integrative Organismal Systems (IOS) Grant 1444612 (Rapid and Targeted Introgression of Traits via Genome Elimination) to L.C.

## Author Contributions

L.C. and E.-H.T. conceived experiments, K.R.A., B.O., E.H.T., M.B., A.K., I.M.H. and L.C. designed experiments, K.R.A., B.O., M.S. and L.C. performed experiments, K.R.A., B.O., M.S., and L.C. performed data analysis, K.R.A., M.L.N., E.H.T., I.M.H. and L.C. interpreted data, B.O., M.S., M.B., and A.K. contributed reagents and materials, and K.R.A. and L.C. wrote the paper with input from all authors.

---

**Supplementary Figure S1.**
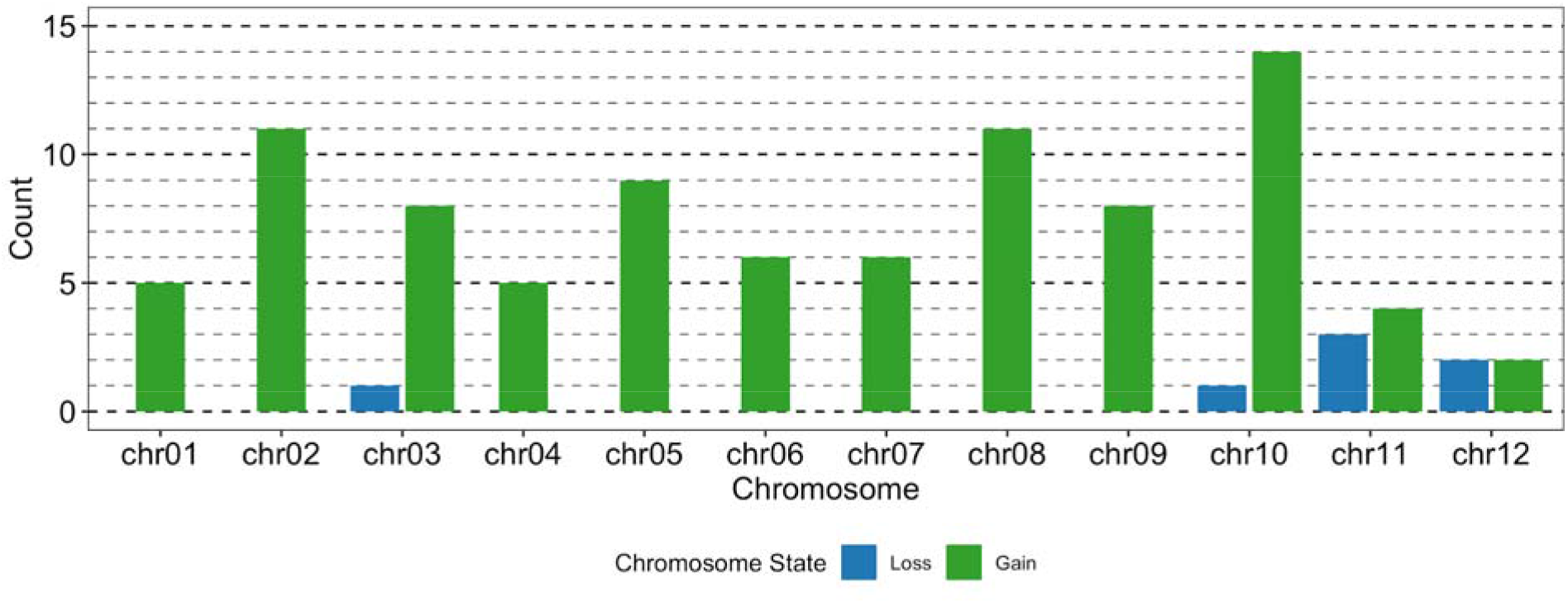
Histogram of aneuploidy by homologous chromosome affected and state loss or gain among 87 potato dihaploids. Instances of chromosomal gain or loss are counted by homologous chromosome and state of loss or gain.

**Supplementary Figure S2.**
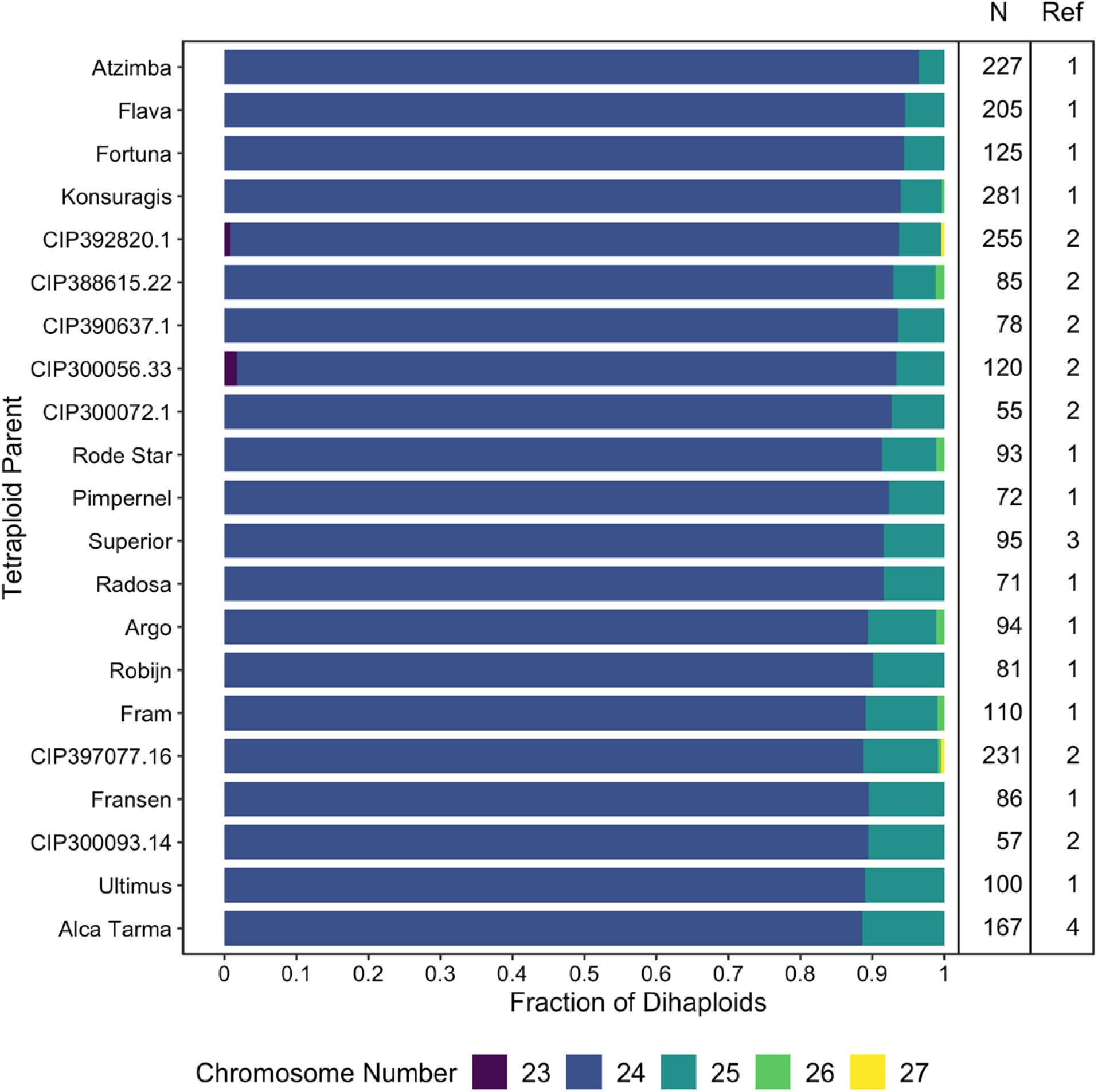
Aneuploidy frequency among putatively uniparental dihaploid potatoes. For each of 21 tetraploid potato clones, the frequency of euploid and various aneuploid karyotypes among extracted dihaploid are reported by chromosome number. The dihaploid population size (N) and corresponding study are listed for reference (Ref) on the right. References: 1) (Wagenvoort and Lange, 1975), 2) this study, 3) (Pham et al., 2019), 4) (Amundson et al., 2020). Only populations with 50 or more dihaploids are shown.

**Supplementary Figure S2.**
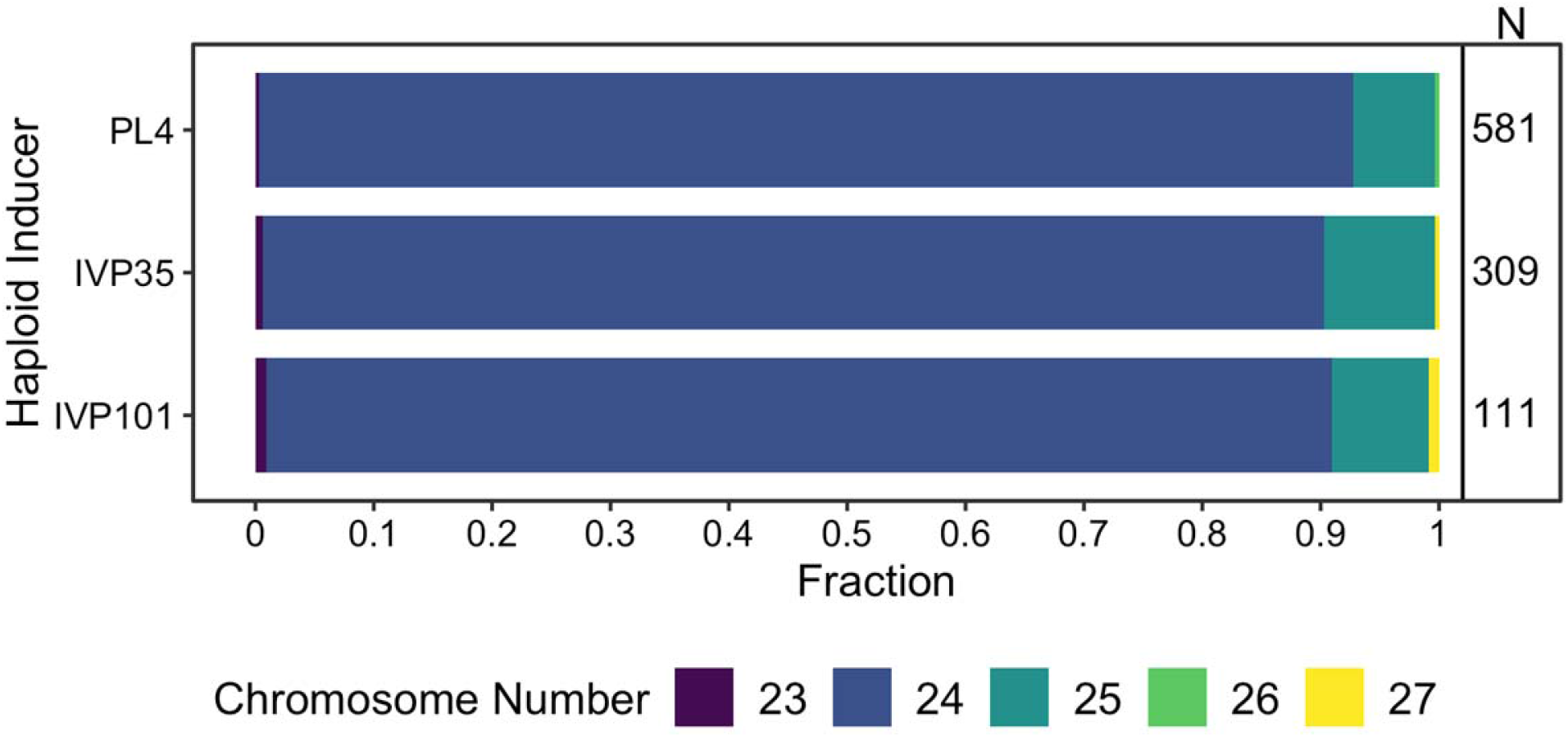
Aneuploidy frequency among putatively uniparental dihaploid potatoes. For each of 3 haploid inducers, the frequency of euploid and various aneuploid karyotypes among extracted dihaploid are reported by chromosome number. The dihaploid population size (N) is listed on the right.

**Supplementary Figure S4.**
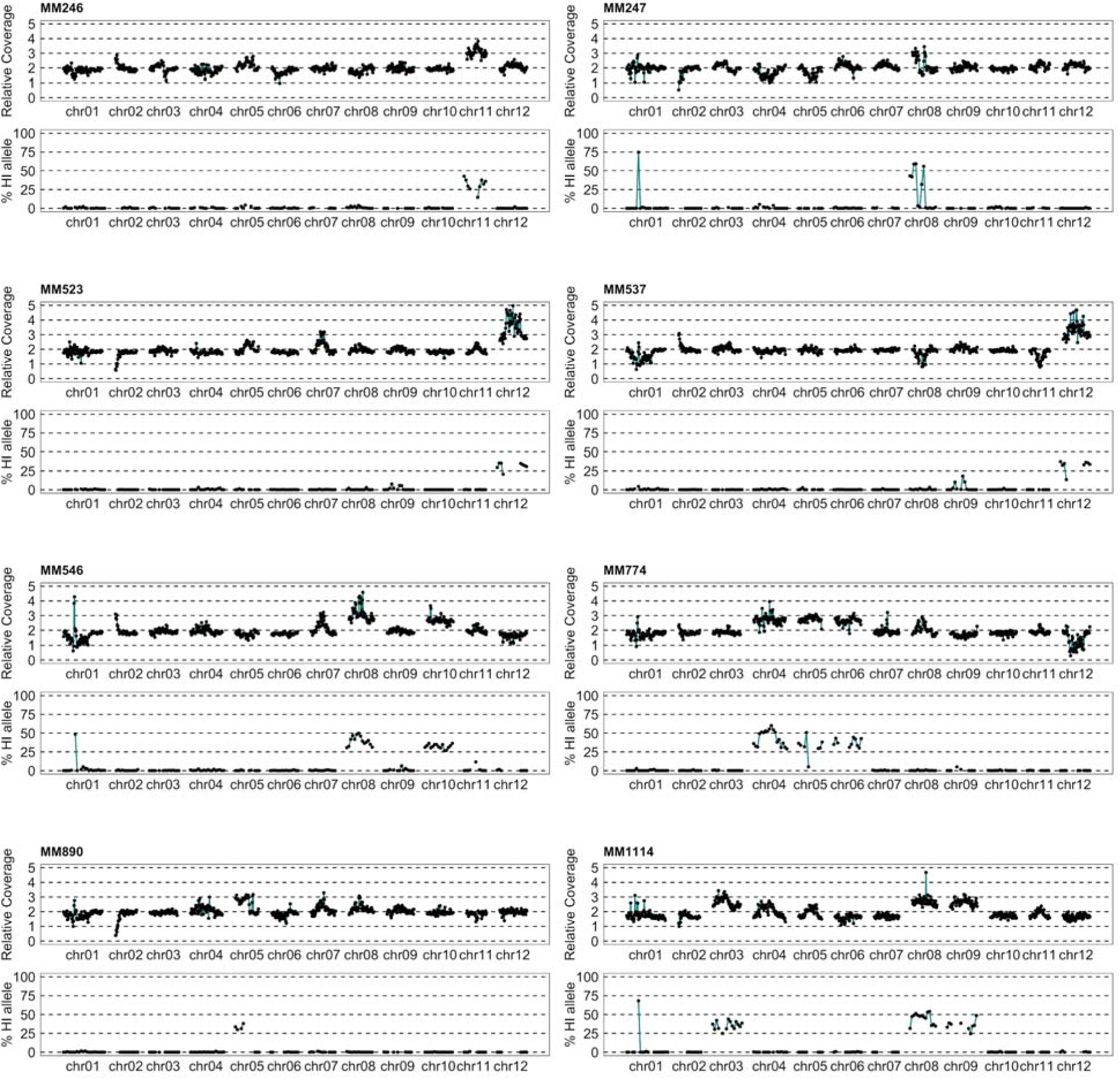
Chromosome dosage and parental allele dosage plots of eight haploid inducer addition dihaploids. Each dihaploid is represented as a pair of vertically stacked plots. The upper plot displayed relative read coverage to each dihaploid’s tetraploid parent. Each point corresponds to the relative coverage of a non-overlapping 1Mb bin of the reference genome. The lower plot displays the percentage of HI-specific allele at all SNP loci identified in non-overlapping 4Mb bins of the reference genome.

**Supplementary Figure S5.**
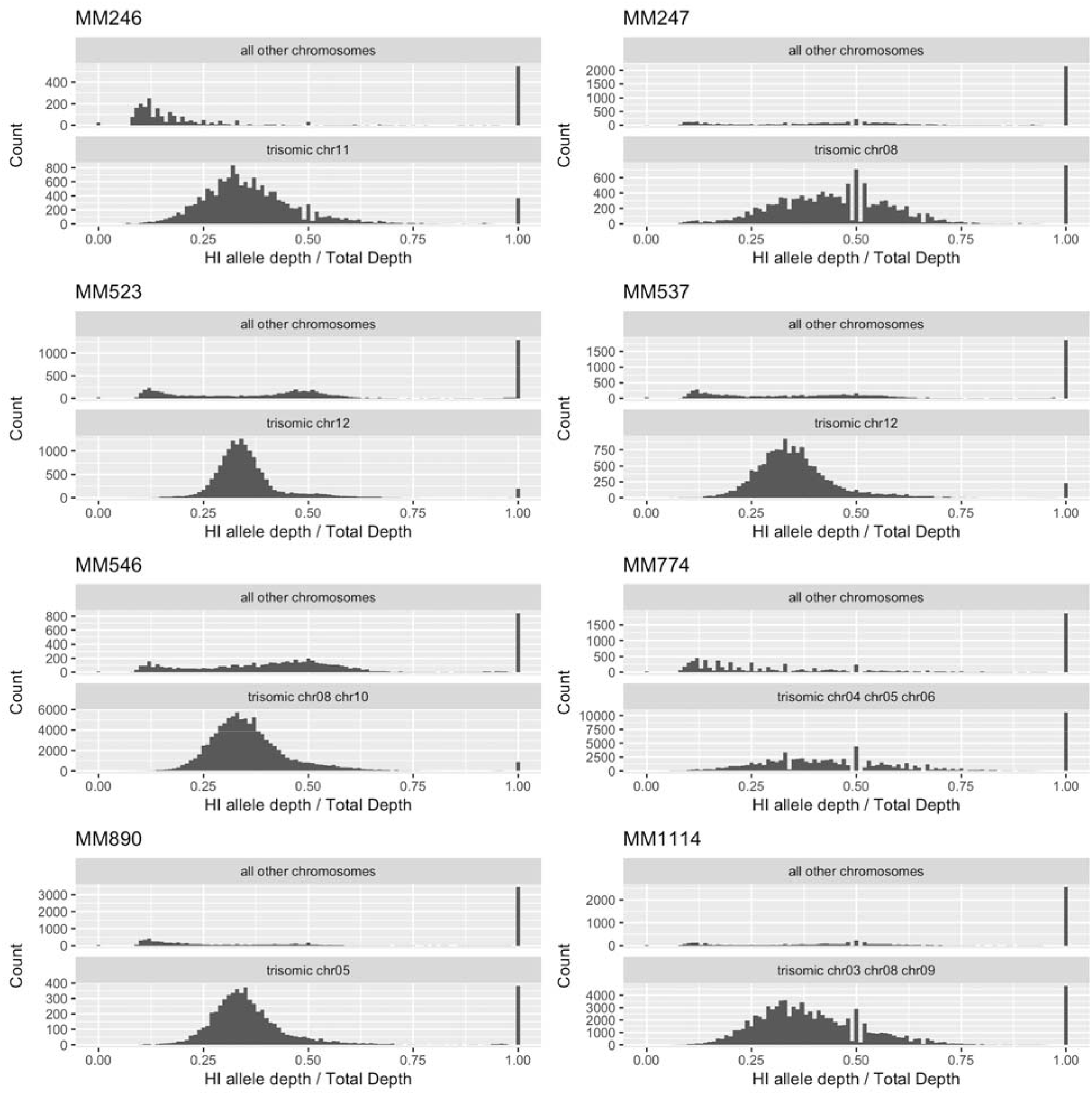
Histograms of haploid inducer allele representation at putative introgression loci in eight HI addition dihaploids.

**Supplementary Figure S6.**
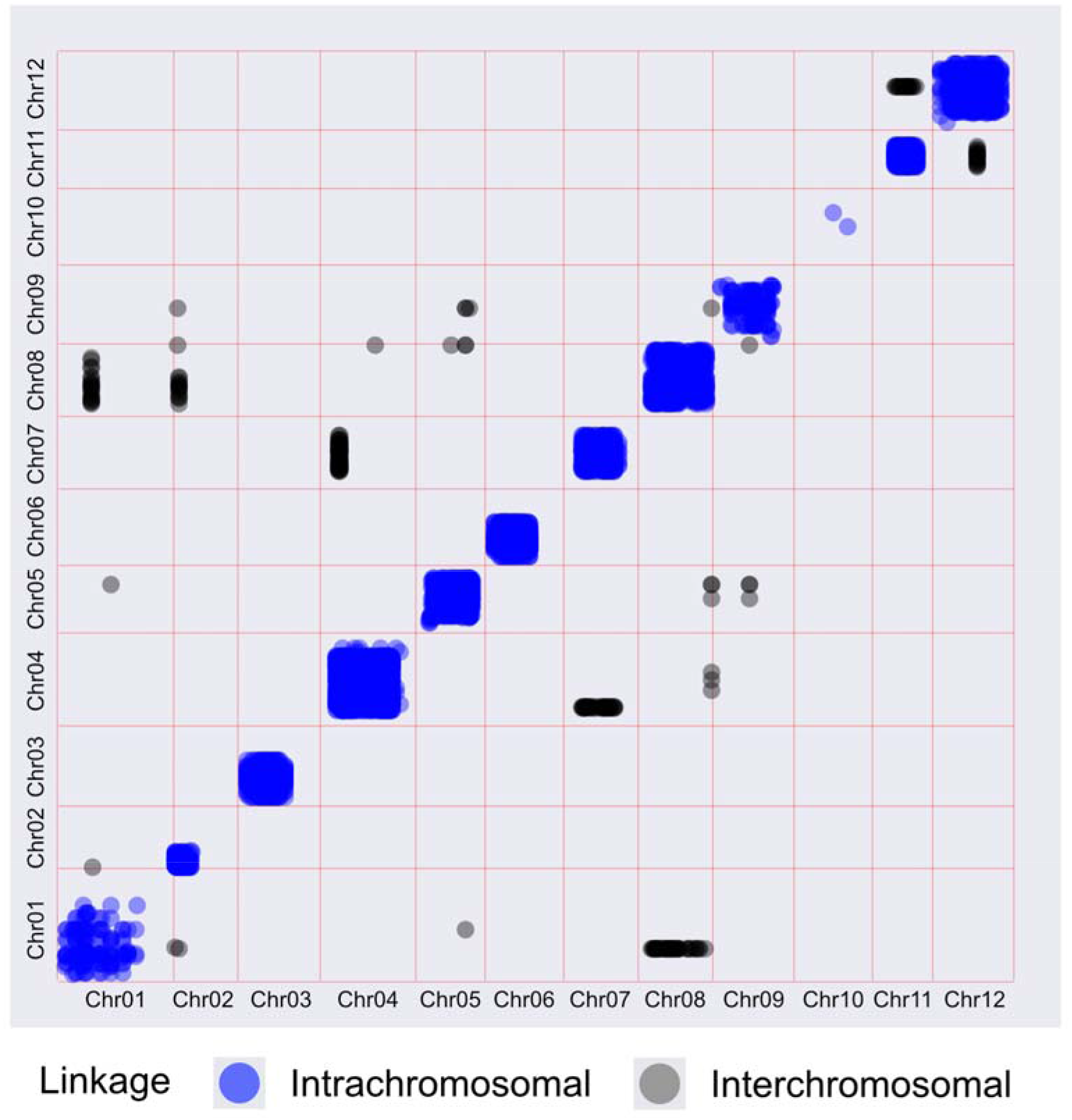
Dosage variation linkage matrix of WA.077 dihaploids. For 231 dihaploids extracted from WA.077, relative coverage values were derived for 1Mb bins and clustered to infer discrete dosage states for each bin. Non-random association between polymorphic bins were assessed by the Fischer Exact test. Blue points: intrachromosomal linkage (False Discovery Rate = 0.05), black points: interchromosomal linkage (FDR = 0.05).

**Supplementary Figure S7.**
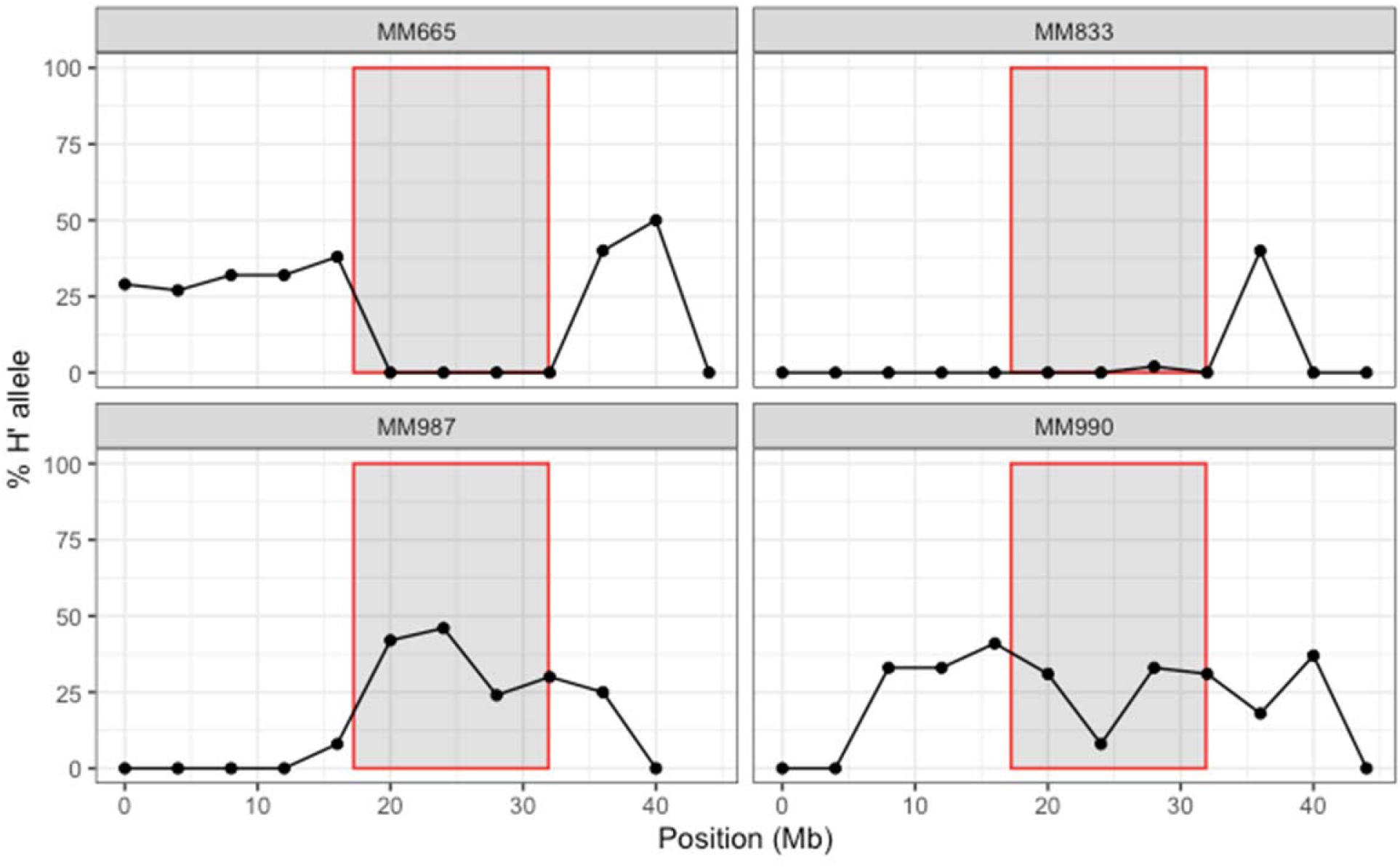
Test of potato haploid inducer (HI) haplotype phasing. From the HI chromosome 11 haplotype extracted from HI addition line MM246 (CIP315048.40), only loci where the phased allele did not match the tetraploid allele were retained. Each panel corresponds to one triploid hybrid. For each hybrid, the percentage of the phased, retained allele (H’ allele) was across SNP loci in non-overlapping 4Mb bins of the reference genome and plotted. The non-recombining region of chromosome 11 reported in (Bourke et al., 2015) is shaded in gray. If the triploid hybrids inherited the same centromere 11 HI haplotype as MM246, approximately 33% H’ allele is expected throughout the entire centromere. If the triploid hybrid inherits the HI centromere 11 haplotype that was not observed in MM246, 0% H’ allele is expected.

**Supplementary Figure S8.**
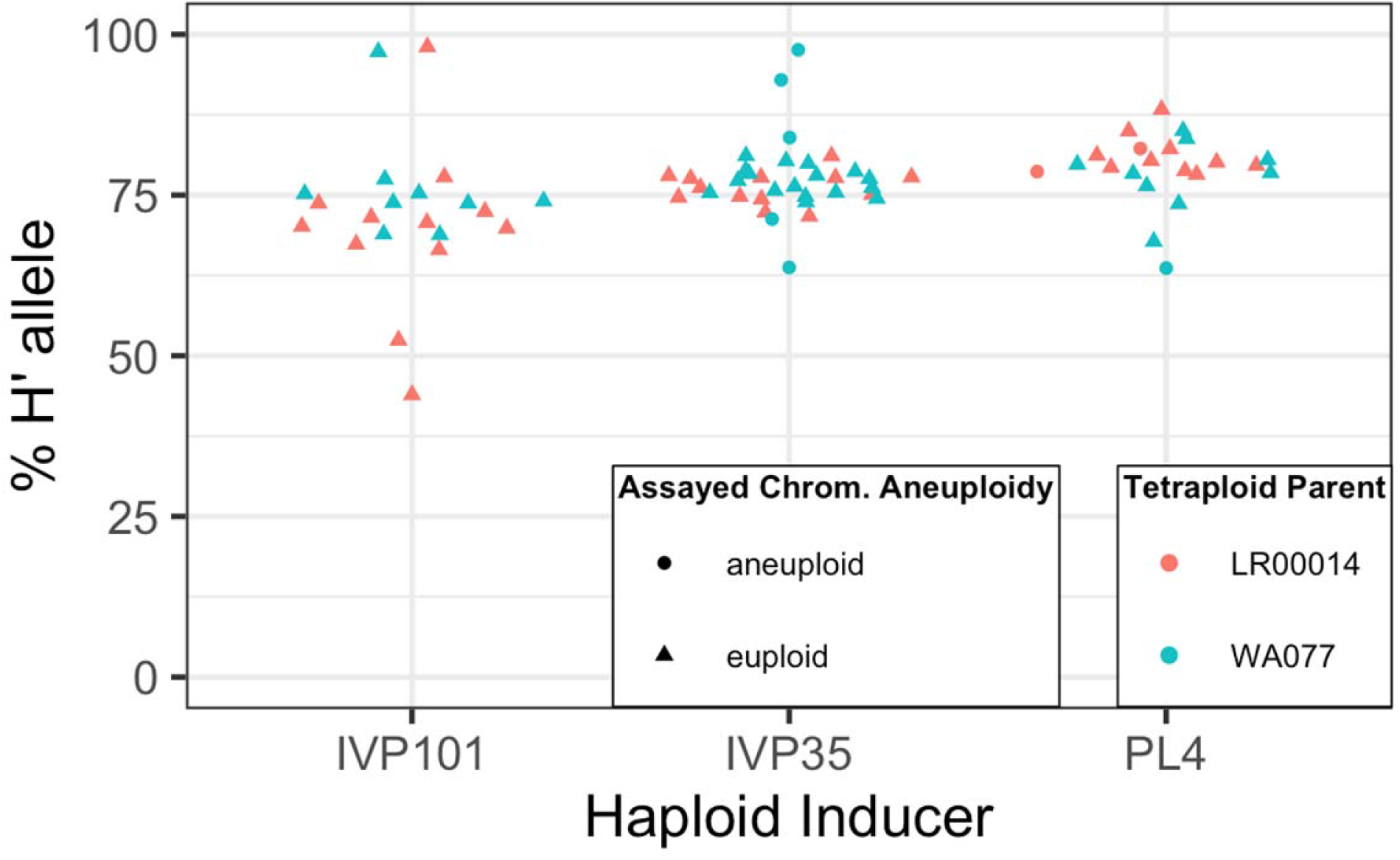
Haploid Inducer (HI) centromeric heterozygosity for tetraploid hybrids of the potato haploid induction cross. Phased HI haplotypes were filtered to retain only loci where the phased HI allele and tetraploid allele were the same, yielding a complementary SNP marker set to the results shown in Fig. 5C. Each point corresponds to the percentage of the phased (H’) allele among reads spanning the non-recombining regions (coordinates from (Bourke et al., 2015)) of a chromosome of one tetraploid hybrid. Chromosome 8 was used to assess IVP101 hybrids. Chromosome 10 was used to assess PL4 hybrids. Chromosome 11 was used to assess IVP35 hybrids. Point shapes indicate whether the assayed chromosome was euploid or euploid.

**Supplementary Figure S9.**
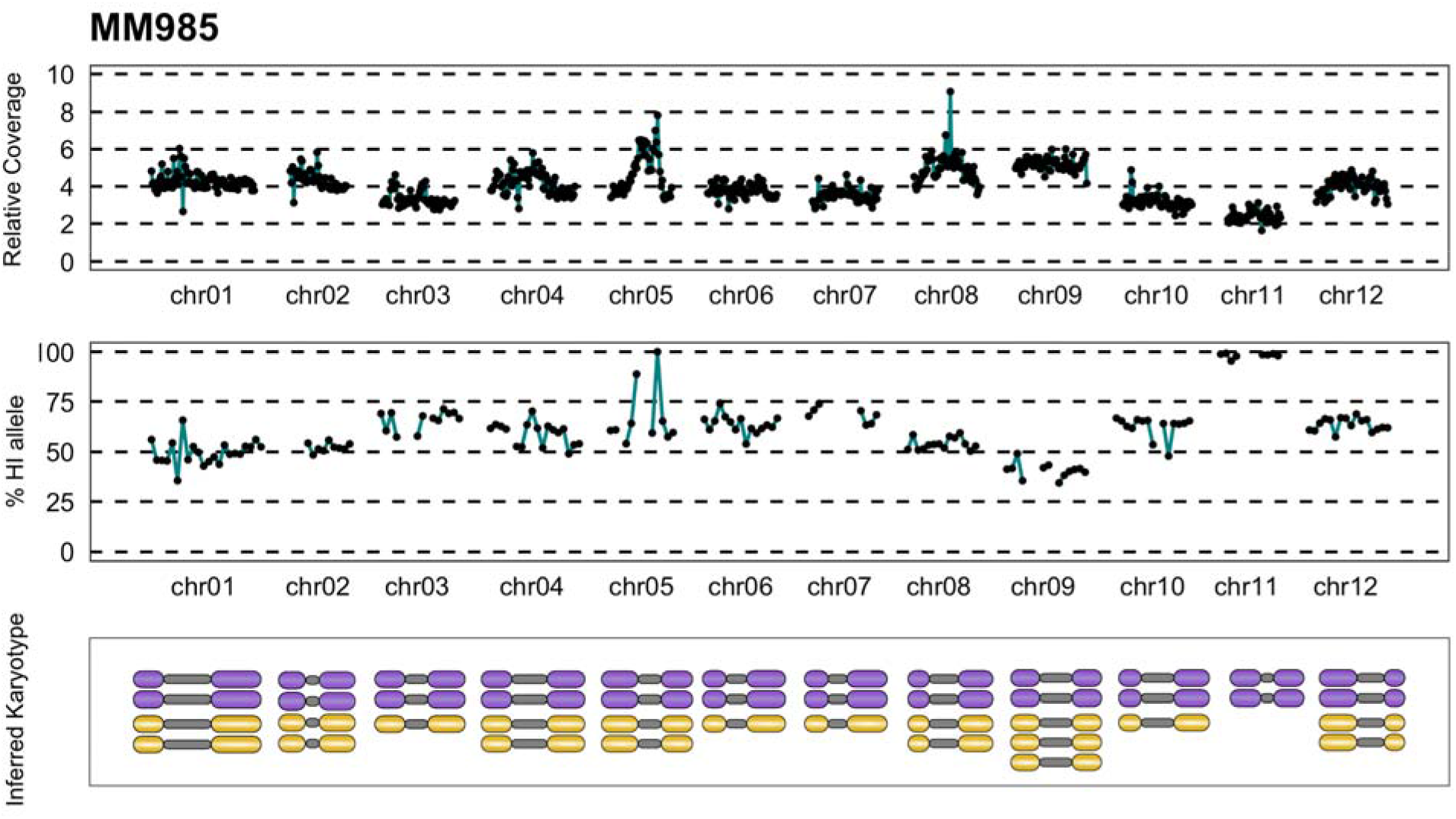
Dosage plot, SNP plot and inferred karyotype of WA.077 x IVP35 chromosome 11 disomic tetraploid potato hybrid.

**Supplementary Table S1.** Genomic content and embryo spot phenotype in progeny of potato haploid induction crosses. For each progeny, ploidy was estimated from flow cytometry or chloroplast counting and presence or absence of the dominant and haploid inducer-specific embryo spot phenotype (Hermsen and Verdenius, 1973) was recorded.

| Ploidy | Without embryo spot | With embryo spot |
| --- | --- | --- |
| 2x | 1041 | 5 |
| 2x-3x | 1 | 0 |
| 3x | 4 | 26 |
| 3x-4x | 1 | 3 |
| 4x | 15 | 114 |
| 5x | 0 | 1 |
| Unknown | 4 | 0 |

**Supplementary Table S2.** Description of 19 tetraploid potato clones used for dihaploid extraction.

| Accession Name | CIP Accession Number | Category |
| --- | --- | --- |
| 458 | CIP391931.1 | Advanced clone |
| 93.003 | CIP390637.1 | Advanced clone |
| Atlantic | CIP800827 | Variety |
| C01.020 | CIP301023.15 | Advanced clone |
| Tacna (C90.170) | CIP390478.9 | Variety |
| C91.640 | CIP388615.22 | Advance clone |
| C92.172 | CIP392780.1 | Advanced clone |
| C93.154 | CIP392820.1 | Advance clone |
| Desiree | CIP800048 | Variety |
| LR00.014 | CIP300056.33 | Advanced clone |
| LR00.022 | CIP300072.1 | Advanced clone |
| LR00.026 | CIP300093.14 | Advance clone |
| LR-93.073 | CIP392822.3 | Advanced clone |
| LRY-3.57 | CIP313047.57 | Advanced clone |
| LRY-21.25 | CIP313065.25 | Advanced clone |
| WA.073 | CIP397099.4 | Advanced clone |
| WA.077 | CIP397077.16 | Advance clone |
| WA.104 | CIP397073.16 | Advanced clone |
| Maria Bonita (Y84.027) | CIP388676.1 | Variety |

## Supplementary Datasets

Dataset S1: Description of plant material used in this study

Dataset S2: Summary of sequencing libraries constructed or analyzed in this study

Both available on Dryad at doi:10.25338/B8JS8D

